# Fine-tuning cellular levels of DprA ensures transformant fitness in the human pathogen *Streptococcus pneumoniae*

**DOI:** 10.1101/314575

**Authors:** Calum Johnston, Isabelle Mortier-Barriere, Vanessa Khemici, Patrice Polard

**Affiliations:** Laboratoire de Microbiologie et Génétique Moléculaires (LMGM), UMR5100, Centre for Integrative Biology (CBI), Centre National de la Recherche Scientifique (CNRS), Toulouse, France; Université de Toulouse, Université Paul Sabatier, Toulouse, France

## Abstract

Natural genetic transformation is a widespread mechanism of bacterial horizontal gene transfer. Transformation involves the internalization of exogenous DNA as single strands, followed by chromosomal integration via homologous recombination, promoting acquisition of new genetic traits. Transformation occurs during a distinct physiological state called competence, during which all proteins required to transform are produced. In the human pathogen *Streptococcus pneumoniae*, competence is controlled by a two-component system ComDE, which is induced by an exported peptide pheromone. DprA is universal among transformable species, strongly and specifically induced during pneumococcal competence, and crucial for pneumococcal transformation. Pneumococcal DprA plays three crucial roles in transformation and competence. Firstly, DprA protects internalized single-stranded (ss) DNA from degradation. Secondly, DprA loads the homologous recombinase RecA onto transforming ssDNA to promote transformation. Finally, DprA interacts with the response regulator ComE to shut-off pneumococcal competence. Pneumococcal shut-off has been linked to physiology, with long growth delays in competent *dprA^-^* cells. Here, we explored the effect of altering the cellular levels of DprA on these three roles. High cellular levels of DprA were not required for the primary role of DprA as a transformation-dedicated recombinase loader or for protection of transforming ssDNA. In contrast, full expression of *dprA* was required for optimal competence shut-off. Full expression of *dprA* was also crucial for transformant fitness. High cellular levels of DprA in competent cells thus ensure the fitness of pneumococcal transformants by promoting competence shut-off. This promotes survival and propagation of transformants, thus maximizing the adaptive potential of this human pathogen.

**Importance:** Transformation is a widespread mechanism of horizontal gene transfer that allows bacteria to acquire new genetic traits by internalizing foreign DNA and integrating it into their chromosomes. Transformation occurs during a transient physiological state called competence. DprA is conserved in transformable species and crucial for the protection and integration of transforming DNA. In the human pathogen *Streptococcus pneumoniae*, DprA is highly abundant and is also crucial for competence shut-off. Here, we show that high DprA expression is not required for transformation. In contrast, full expression of *dprA* was required for competence shut-off and transformant fitness. These findings thus link high cellular levels of DprA to survival and propagation of pneumococcal transformants, maximizing the adaptive potential of this human pathogen.

## Introduction

Natural genetic transformation is a mechanism of horizontal gene transfer which is widespread in the bacterial kingdom 1. This process, entirely encoded by the recipient cell, involves the capture and internalization of exogenous DNA followed by integration into the recipient chromosome by homologous recombination 1. Transformation thus allows acquisition of new genetic traits and promotes the spread of antibiotic resistance and vaccine escape 2. In the human pathogen *Streptococcus pneumoniae* (the pneumococcus), transformation occurs during a short window called competence, which occurs during early exponential growth 3. Pneumococcal competence is a transient physiological state during which over 100 genes are specifically expressed 4, 5. Around 20 of these are involved in transformation 6, with the products of these genes collectively defined as the transformasome 7.

In *S. pneumoniae*, the transformation process begins with the transformation pilus 8, 9, encoded by the *comG* operon 10, 11, which captures exogenous DNA 8. This DNA is then accessed by the DNA receptor ComEA, and the endonuclease EndA, which degrades one strand 12, 13, allowing the remaining strand to be internalized through the ComEC transformation pore, mediated by the ATP-dependent DNA translocase ComFA 14. Upon internalization, transforming ssDNA initially interacts with three transformasome proteins, DprA, RecA and SsbB. The transformation-dedicated recombination mediator protein (RMP) DprA can bind to ssDNA and load the homologous recombinase RecA to promote homologous recombination 15, 16. Both DprA and RecA protect transforming ssDNA from degradation 17. Once RecA is loaded onto ssDNA, it polymerizes and promotes chromosomal integration by homologous recombination. Interestingly, DprA is produced at high levels, with over 8,000 molecules present in each competent cell 18. This is in stark contrast to expression levels of other RMPs such as RecFOR and RecBCD, with less than 100 molecules generally thought to be present per cell 19. Transforming ssDNA can also be coated with the transformation-dedicated single-stranded DNA-binding protein SsbB 20, which generates a reservoir of DNA for transformation, facilitating multiple transformation events 21.

Competence is a transient phenotype and uncontrolled induction was observed to strongly impact pneumococcal growth in *in vitro* planktonic culture 18. Induction and shut-off of competence are thus tightly regulated. Firstly, induction is controlled by an autocatalytic feedback loop involving the genes *comABCDE*. The *comC* gene encodes the peptide pheromone CSP (Competence Stimulating Peptide), which is exported and matured by the dedicated ComAB ABC transporter 22. Outside the cell, CSP induces phosphorylation of the membrane-bound histidine kinase ComD, which in turn phosphorylates the intracellular response regulator ComE. ComE~P induces the expression of the *comAB* and *comCDE* operons, generating an autocatalytic feedback loop, as well as other ‘early’ competence (*com*) genes. The early *com* genes *comX1* and *comX2* encode for an alternative sigma factor (σ^X^), which induces the ‘late’ *com* genes, including those of the transformasome 7, 23. Secondly, competence shut-off is mediated by DprA, which interacts strongly with ComE~P to either sequester or dephosphorylate it, promoting the shut-off of competence 18. Inactivation of *dprA* was shown to be highly detrimental for development of pneumococcal infection, dependent on the ability of cells to develop competence 24. This is probably linked to the large growth defect observed in competent *dprA^-^* cells grown *in vitro*, which is due to uncontrolled competence induction 18. Thus, DprA plays key roles in transformation and competence (Figure 1A), and the DprA-mediated shut-off of pneumococcal competence is key for fitness. This study aimed to investigate how these two key activities are balanced in pneumococcal cells.

**Figure 1:**
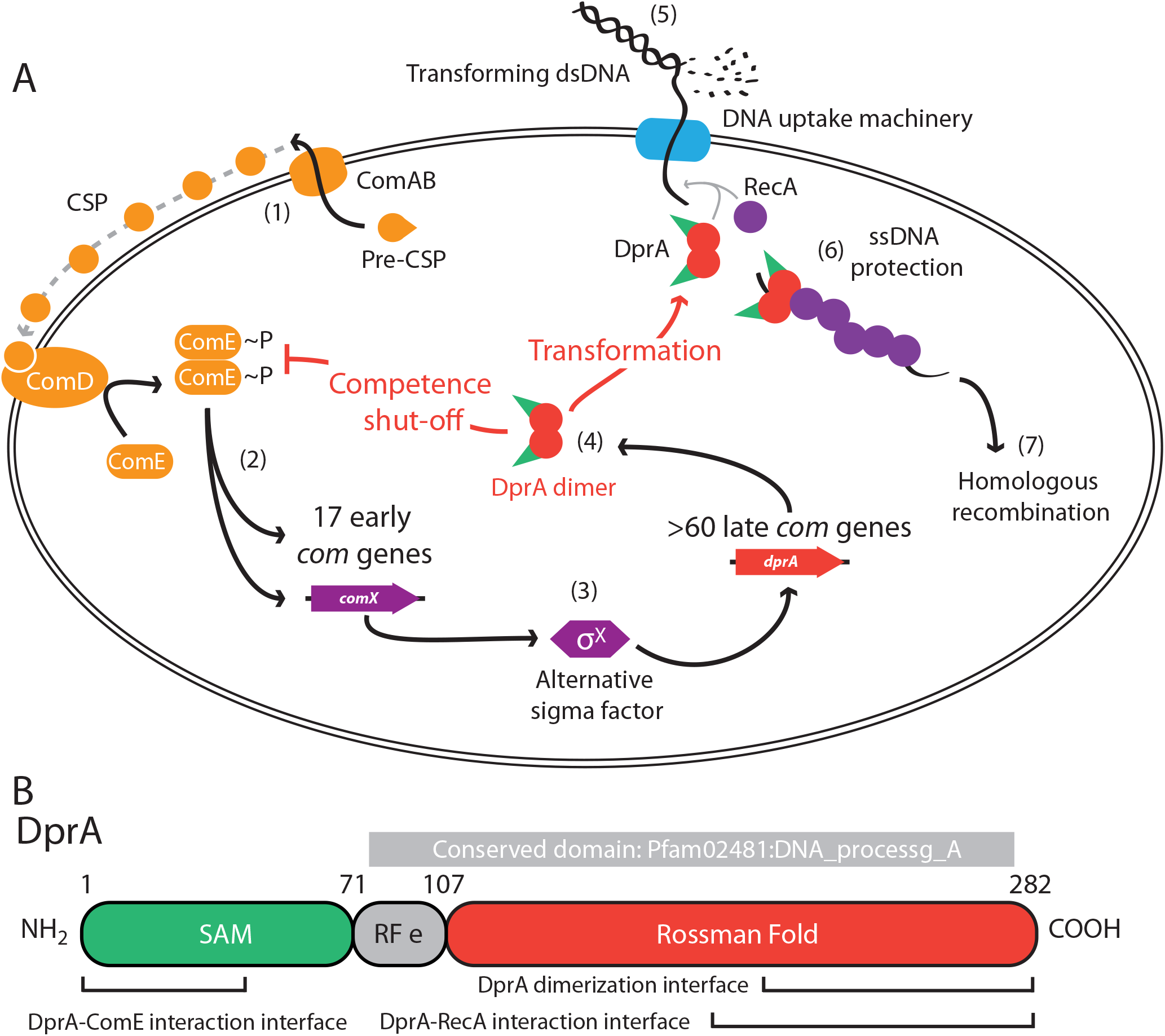
DprA in competence and transformation. (A) 1 Pre-CSP, encoded by the *comC* gene, is exported and matured by the ComAB transporter, and then interacts with the histidine kinase ComD, stimulating its phosphorylation. 2 ComD~P transphosphorylates the response regulator ComE, which then stimulates the expression of 17 early *com* genes, including two copies of *comX*. 3 These encode an alternative sigma factor σ^X^, which induces expression of over 60 late *com* genes including *dprA*. 4 DprA dimers load RecA onto ssDNA to mediate transformation and interact with ComE~P to shut-off competence. 5 Transforming DNA is internalized in single strand form via the DNA uptake machinery. 6 DprA and RecA interact with transforming ssDNA, protecting it from degradation. 7 RecA mediates homologous recombination between transforming ssDNA and the chromosome to promote transformation. (B) Linear representation of DprA with the limits of Pfam02481 (in gray) and structural domains indicated. Interfaces of dimerization and interaction with RecA and ComE are indicated and represent the regions within which the majority of point mutations affecting interaction were identified 16, 18.

Pneumococcal DprA is made up of two domains (Figure 1B), an N-terminal sterile alpha motif (SAM) domain and a C-terminal extended Rossman fold (eRF) 16. The interaction of DprA with its partner proteins involves distinct domains, with interaction with ComE~P occurring via the SAM domain 18 and dimerization and interaction with RecA involving the eRF 16. DprA is conserved among all transformable species 1, and absence of pneumococcal DprA results in almost complete loss of transformation 17. In *Bacillus subtilis*, DprA also loads RecA onto transforming ssDNA 25, although inactivation of *dprA* results in a more modest ~50-100-fold loss of transformation efficiency 26. DprA also plays a role in transformation, presumably as a RMP, in *Haemophilus influenzae* 27, *Neisseria gonorrhoeae* 28, *Neisseria meningitidis* 29 *Campylobacter jejuni* 30 and *Helicobacter pylori* 31, 32. Although the mechanism of transformation remains broadly conserved across transformable bacteria, the regulation of competence is very diverse 1. Since pneumococcal competence shut-off is dependent on interaction between DprA and ComE~P, the role of DprA in competence shut-off may be limited to streptococci which control competence via ComDE 18.

In this study, we explore the effect of altering the cellular levels of DprA on the roles of DprA in transformation and competence shut-off. By controlling cellular DprA levels using an inducible expression platform, we show that even a ten-fold reduction in cellular DprA barely affects transformation. In contrast, optimal competence shut-off requires full *dprA* expression. The high cellular level of DprA is thus coupled to its distinct role in competence shut-off and as a result may only be conserved among streptococci that control competence via ComDE. In addition, high cellular levels of DprA are crucial for the fitness of pneumococcal transformants. Together, these results directly link high cellular levels of DprA to survival and propagation of pneumococcal transformants, optimizing the adaptive potential of this human pathogen.

## Results

### Controlling cellular levels of DprA using CEP_*lac*_

The aim of this study was to explore the effect of varying the cellular levels of DprA on the roles of this protein in pneumococcal transformation and competence shut-off. In order to achieve this, a platform allowing expression of a desired gene from the P_*lac*_ promoter 33 was constructed and validated using the *luc* gene 34 as described in the Supplementary Information (Figure S1). To explore the effect of varying cellular DprA levels on its roles in competence shut-off and transformation, the *dprA* gene was placed in the CEP platform 35 controlled by the P_*lac*_ promoter to generate CEP_*lac*_*-dprA* (Figure 2A), before inactivation of native *dprA* using a spectinomycin resistance cassette (*dprA::spc^21^*^C^, 18) Western blots using α-DprA antibodies show that expression levels of *dprA* can be finely tuned by varying IPTG concentration (Figure 2B). To determine the number of DprA molecules per cell in each IPTG concentration, quantitative Western blots were carried out as previously described 18. Results show that ~8,250 wildtype DprA molecules are produced per competent cell, in agreement with previous findings 18, validating our quantifications (Figure 2C). In contrast, maximal induction of CEP_*lac*_*-dprA* generated ~6,500 DprA molecules per cell, with a decreasing IPTG concentration resulting in a decreasing number of DprA molecules per cell, with only ~40 molecules observed in absence of IPTG (Figure 2C). This strain thus provided the opportunity to test how altering cellular concentrations of DprA affects transformation and competence shut-off.

**Figure 2:**
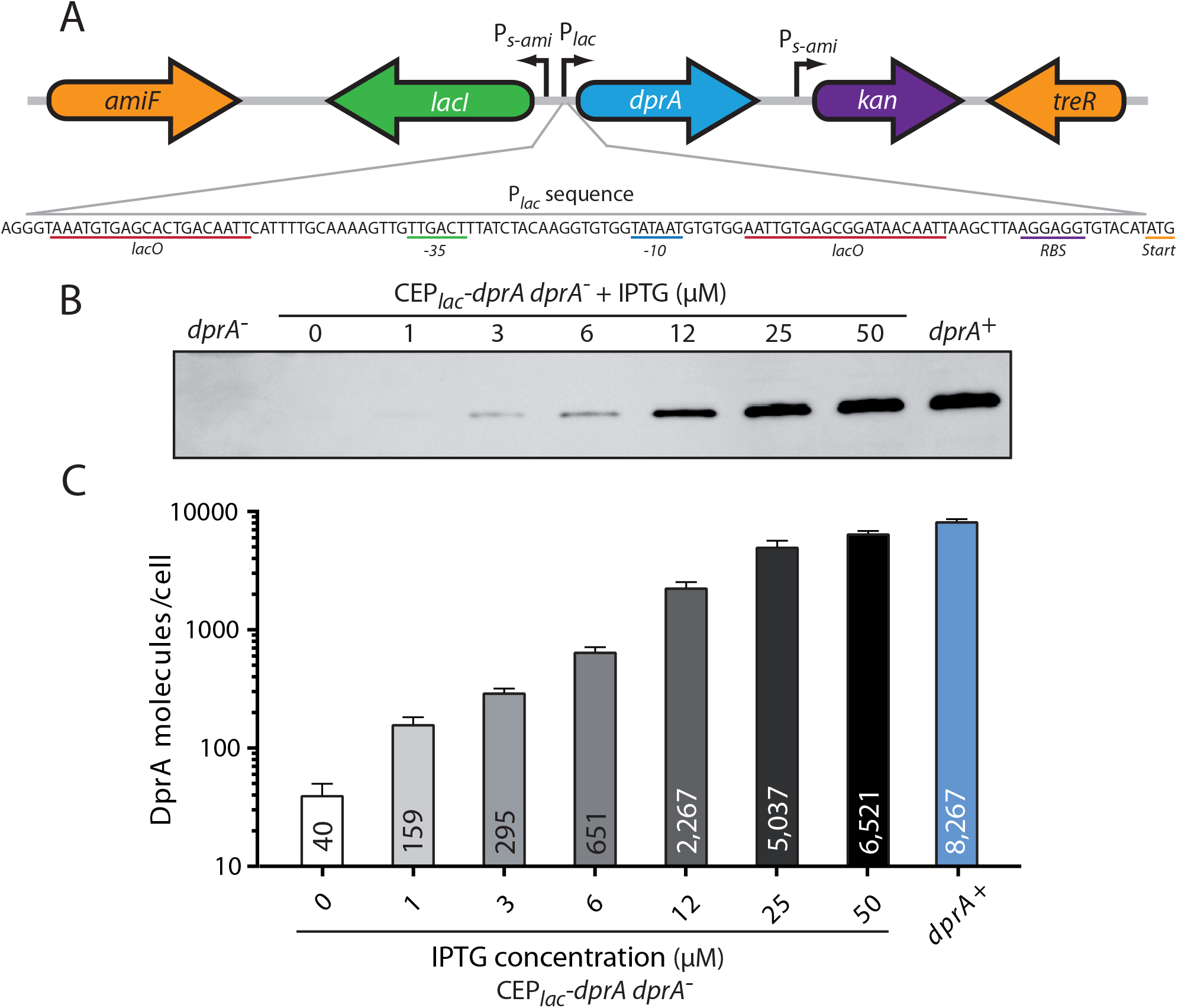
Quantification of DprA produced from CEP_*lac*_*-dprA*. (A) Genetic organization of CEP_*lac*_*-dprA* expression platform. *amiF* and *treR* are pneumococcal genes between which the platform is inserted. The *kan* gene is constitutively expressed via the P_*s-ami*_ promoter and confers kanamycin resistance. The *lacI* repressor gene is also expressed via a distinct P_*s-ami*_ promoter. The *dprA* gene is expressed via P_*lac*_, the sequence of which is shown. RBS, ribosome-binding site; Start, start codon (ATG). (B) Western blot comparing expression levels of CEP_*lac*_*-dprA* in varying IPTG concentrations to those of wildtype and *dprA^-^* cells. (C) Quantified numbers of DprA molecules produced per competent cell for each concentration of IPTG. Results represent averages of triplicate repeats of Western blots compared to gradients of purified DprA. Strain used: CEP_*lac*_*-dprA*, *dprA^-^*, R3833.

### Optimal transformation requires few DprA molecules per competent cell

The conserved role of DprA is as a RMP, to load RecA onto transforming ssDNA during transformation 15. To investigate whether reducing cellular DprA levels impacted pneumococcal transformation, the transformation efficiency of CEP_*lac*_*-dprA*, *dprA^-^* cells in varying concentrations of IPTG was determined. To achieve this, CEP_*lac*_*-dprA*, *dprA^-^* cells were transformed with either saturating concentrations of genomic DNA possessing a point mutation conferring streptomycin resistance (*rpsL41*), or non-saturating concentrations of a PCR fragment possessing the same point mutation, where on average one DNA molecule is present per three competent cells. In saturating DNA concentrations (Figure 3A), there was no significant difference between transformation efficiencies in presence of 12-50 μM IPTG (2,200-6,500 DprA molecules per competent cell; Figure 2A). With 650 DprA molecules per competent cell, there is a modest but significant three-fold drop in transformation efficiency, followed by a rapid linear decline in transformation efficiency below this point. Indeed, in the absence of IPTG, where only ~40 DprA molecules are produced per competent cell, transformation efficiency drops ~240-fold. This remains above the >5-log drop observed in *dprA^-^* cells 15, showing that even 40 molecules of DprA are capable of ensuring transformation. Taken together, these results show that full transformation efficiency is maintained in presence of ~650-2,250 molecules of DprA, but that below this point, the cellular levels of DprA fall below those necessary to stimulate optimal transformation. In non-saturating DNA concentrations, the profile of efficiency was similar, although there was no significant difference between ~6,500 and ~650 molecules of DprA (Figure 3B). This result shows that even when transforming DNA is limiting, ~300 DprA molecules per cell (3 μM IPTG) are not sufficient to ensure optimal transformation efficiency. Fold decreases in transformation efficiency taken for each cellular level of DprA compared to the highest expression condition are directly comparable between excess and limiting transforming DNA conditions (Figure 3C). This demonstrates that loss of transformation efficiency directly correlates to cellular DprA levels independent of the concentration of transforming DNA present. Taken together, these results show that whether the transforming DNA is in excess or limiting, between ~650 and ~2,250 molecules of DprA per competent cell are required to ensure maximal transformation efficiency.

**Figure 3:**
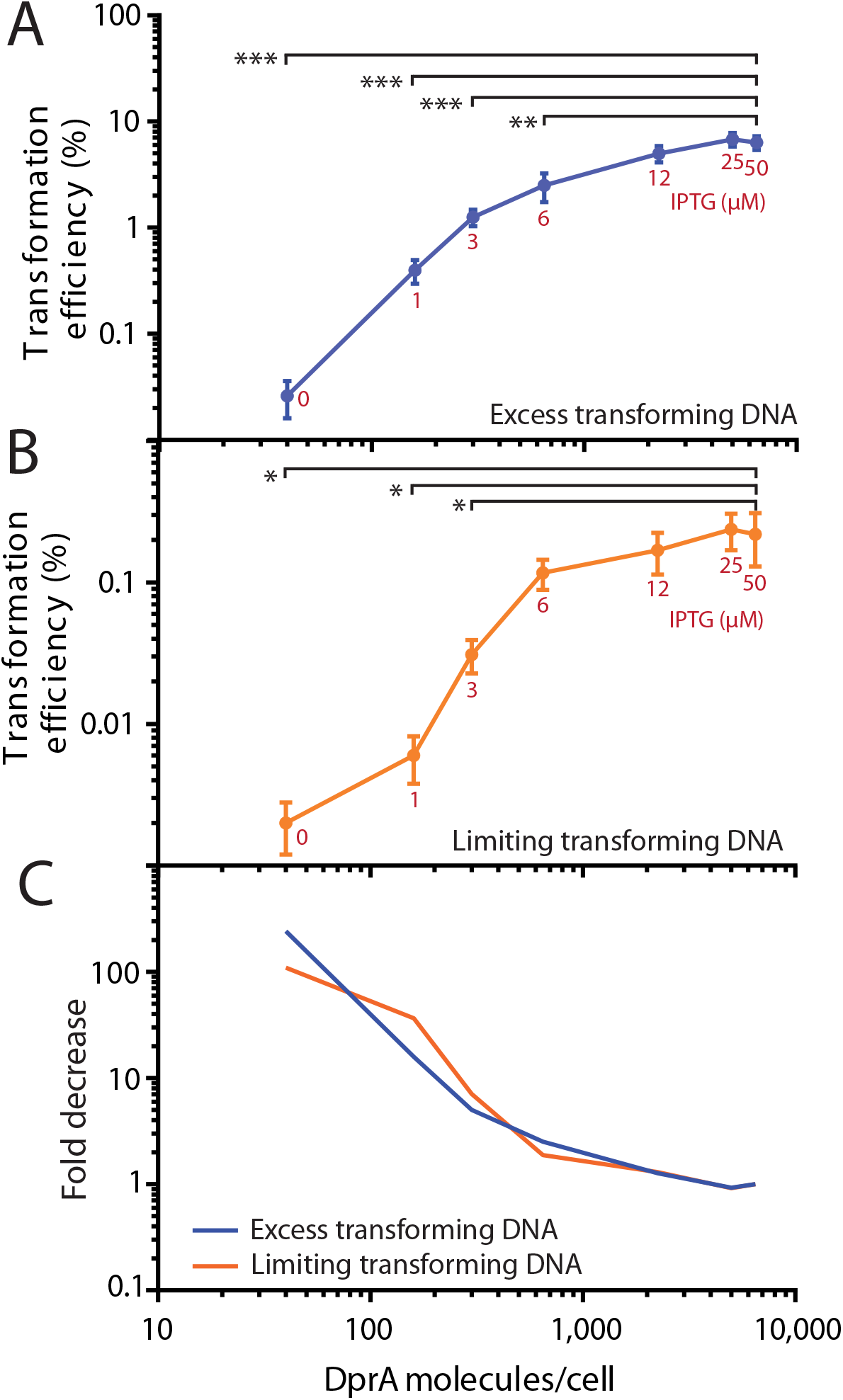
The effect of varying IPTG concentration on transformation efficiency in CEP_*lac*_*-dprA*, *dprA^-^*. (A) Transformation efficiencies of CEP_*lac*_*-dprA*, dprA^-^ strain in varying IPTG concentrations when transformed with excess transforming DNA. Results represent averages of triplicate repeats. **, p<0,01; ***, p<0,001. (B) Transformation efficiencies of CEP_*lac*_*-dprA*, dprA^-^ strain in varying IPTG concentrations when transformed with limiting transforming DNA (~1 DNA molecule per 3 competent cells). Results represent averages of triplicate repeats. *, p<0,05. (C) Fold decrease of transformation efficiencies compared to condition with highest expression of DprA (50μM IPTG). Blue line, excess DNA; orange line, limiting DNA. Strain used: CEP_*lac*_*-dprA*, *dprA^-^*, R3833.

### DprA-mediated protection of transforming ssDNA correlates with transformation efficiency

DprA plays two roles that directly affect transformation, as a recombination mediator protein and as a protector of transforming ssDNA. Since both DprA and RecA are crucial for protection of transforming ssDNA, it is possible that protection and recombination are one and the same, in that the DprA-mediated loading of RecA onto transforming ssDNA, and subsequent polymerization, itself protects the DNA. To shed light on this, the protection of transforming ssDNA was compared in competent cells expressing varying levels of DprA. To achieve this, competent cells were transformed with radio-labelled PCR fragments and cellular DNA recovered to determine the presence of internalized radiolabeled ssDNA, resolved and visualized by native agarose gel electrophoresis. When comparing wildtype and *dprA^-^* cells, a large smear of protected ssDNA was observed in wildtype at both timepoints, while no signal was detected for *dprA^-^* cells (Figure 4A), as previously reported 17, 36, indicating rapid degradation of transforming ssDNA after internalization. For CEP_*lac*_*-dprA*, *dprA^-^* cells, 25 μM IPTG showed ssDNA protection similar to wild-type, followed by a loss of protection correlating with decreasing IPTG concentration (Figure 4A). Density plots of the ssDNA smears in each lane (Figure 4BC) and calculation of the area under these plots (Figure 4D) allowed quantification of ssDNA protection and showed that below 6μM of IPTG (~650 DprA molecules/cell), almost no protection of transforming ssDNA was observed (Figure 4BCD), correlating with the rapid loss of transformation efficiency observed in these conditions (Figure 3AB). Taken together, these results show a strong correlation between protection of ssDNA and transformation efficiency. This strongly suggests that ssDNA protection mediated by DprA limits transformation efficiency.

**Figure 4:**
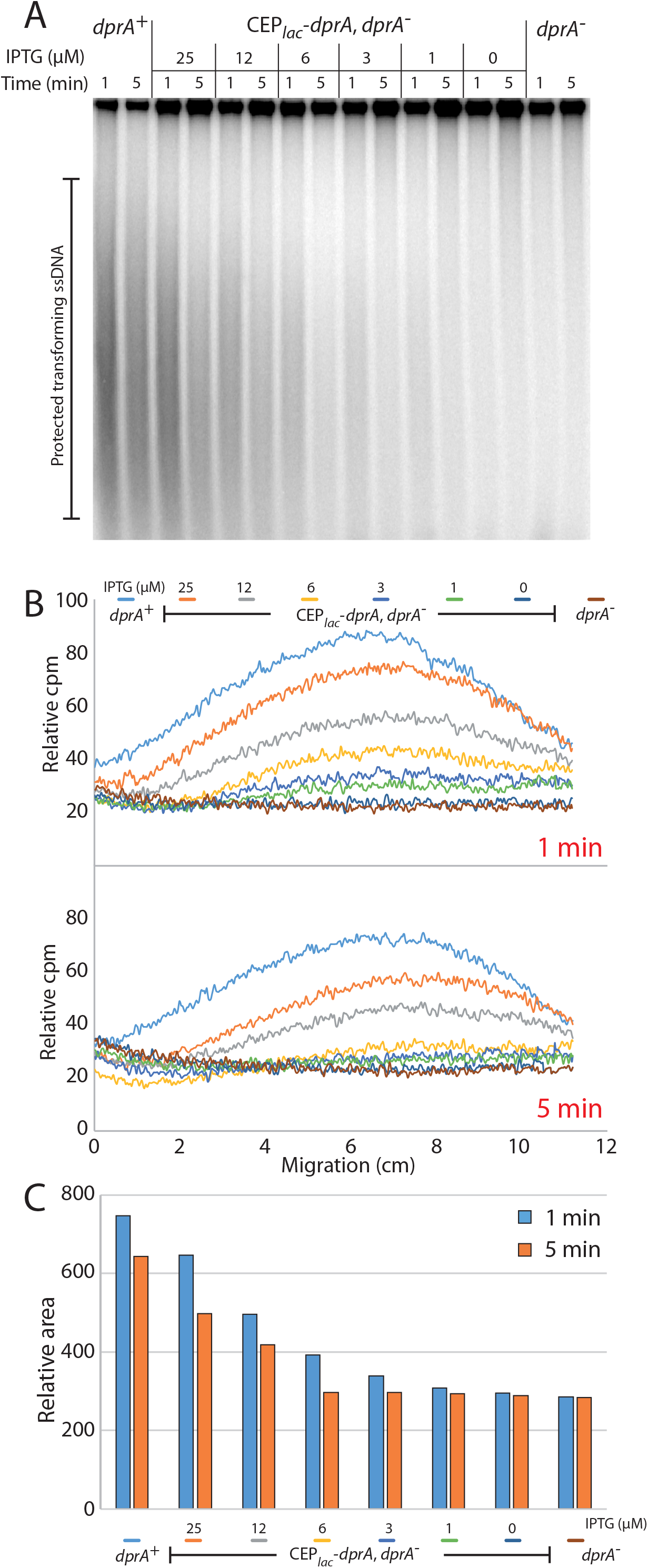
The effect of varying IPTG concentration on ssDNA protection in CEP_*lac*_*-dprA*, *dprA^-^*. (A) Electrophoregram of agarose gel of DNA extracted from cells transformed with radio-labelled DNA in varying IPTG concentrations. Top bands represent chromosomal DNA, and smears represent radio-labelled transforming ssDNA. Time represents time after blockage of DNA uptake by DNAse addition. (B) Density plots of different lanes expressed in relative counts per million. (C) Plots of the relative areas underneath the density curves calculated in panel B. Strains used: *dprA^+^*, R3584; CEP_*lac*_*-dprA*, *dprA^-^*, R3833; *dprA^-^*, R3587.

### Protection of transforming ssDNA depends on DprA-mediated RecA loading on ssDNA

Since transforming ssDNA protection is dependent on not only DprA but also RecA 17, we hypothesized that DprA-mediated RecA-loading protected target ssDNA from degradation. To test this hypothesis, the protection of transforming ssDNA was investigated in a strain possessing a DprA mutant specifically affected for its interaction with RecA (DprA^QNQ^, 16). In addition, the ability of DprA to dimerize is as important for transformation as its ability to interact with RecA 16. The stability of transforming ssDNA in a DprA dimerization mutant (DprA^AR^, 16), which is able to interact with RecA but interacts poorly with ssDNA, was thus also analyzed. These two DprA mutants were incapable of protecting transforming ssDNA (Figure 5A). Density plots of each lane confirmed that these two mutants do not detectably protect ssDNA better than cells lacking *dprA* (Figure 5BC). Thus, the dimerization of DprA and its interaction with RecA are both crucial for transformation 16 but also for protection of transforming ssDNA. Taken together with the fact that transformation and ssDNA protection require similar cellular levels of DprA, these results strongly suggest that the actions of DprA in both of these processes stem from a single mechanism: the DprA-mediated loading of RecA onto transforming ssDNA.

**Figure 5:**
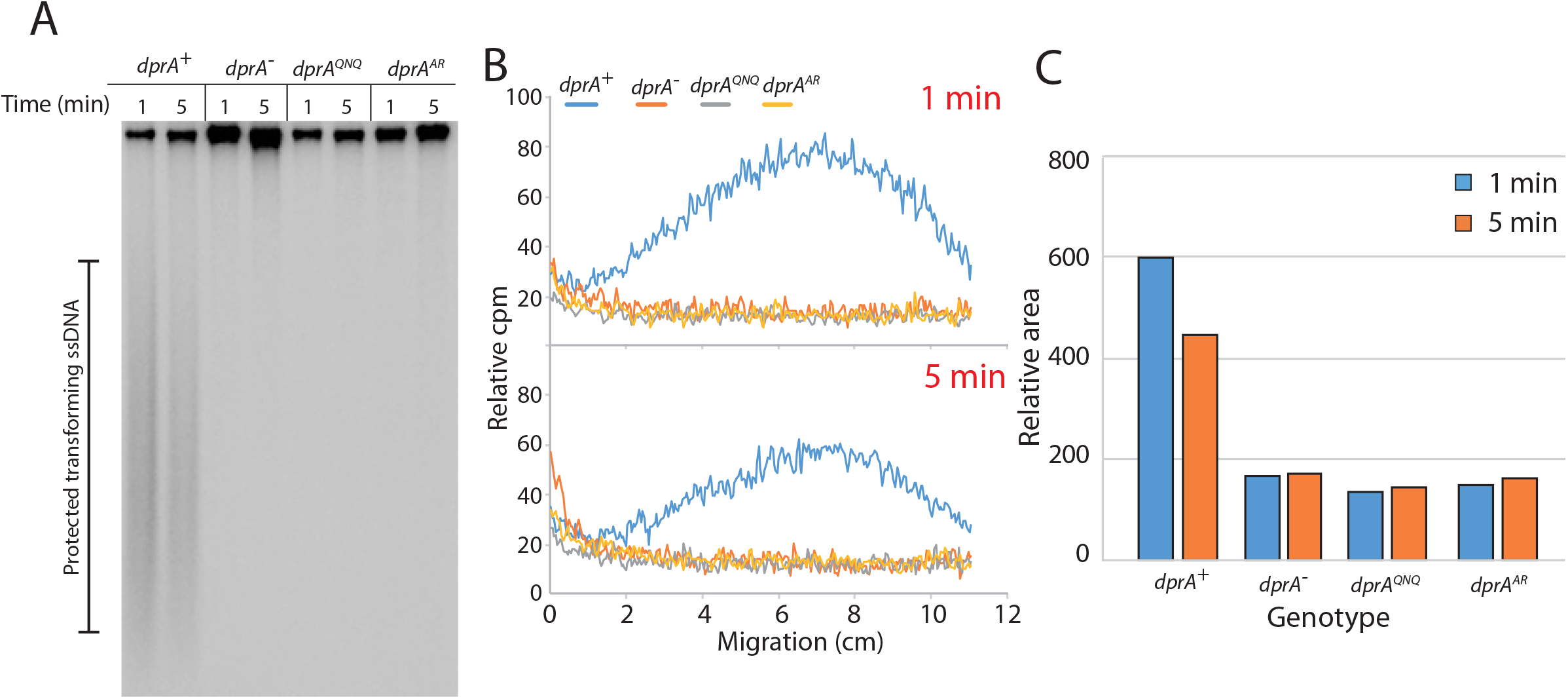
Transforming ssDNA protection in DprA^AR^ and DprA^QNQ^ mutants. (A) Electrophoregram of agarose gel of DNA extracted from cells transformed with radio-labelled DNA. Top bands represent chromosomal DNA, and smears represent radio-labelled transforming ssDNA. Time represents time after blockage of DNA uptake by DNAse. (B) Density plots of different lanes expressed in relative counts per million. (C) Plot of the relative areas underneath the density curves calculated in panel B. Strains used: *dprA^+^*, R1818; *dprA^-^*, R2017; *dprA^QNQ^*, R2830; *dprA^AR^*, R3108.

### Optimal competence shut-off requires full expression of *dprA*

The master regulator of competence ComE is produced at over 80,000 molecules per competent cell 37, although it is unclear what proportion of this is phosphorylated. In addition, previous experiments showed that full expression of *dprA* was not required for optimal transformation (Figure 3). We next explored whether the high levels of DprA expression during competence were necessary for its interaction with ComE~P to mediate competence shut-off. To determine the impact of reducing cellular levels of DprA on competence shut-off, a transcriptional fusion consisting of the promoter of the late *com* gene *ssbB* and the *luc* reporter gene 38, which allows tracking of competence in live cells in real time, was inserted into the CEP_*lac*_*-dprA*, *dprA*^-^ strain. The shut-off of competence was determined in this strain in the same IPTG gradient as used above, compared to wildtype and *dprA^-^* strains. Competence was rapidly induced in wildtype cells, reaching a peak around 20 minutes post-induction, followed by shut-off (Figure 6A). In the absence of *dprA*, no competence shut-off was observed (Figure 6A), in line with previous results 18. In the CEP_*lac*_*-dprA*, *dprA*^-^, *ssbB-luc* strain, the peak of competence occurs slightly later than in the wildtype strain (Figure 6B), possibly due to the ectopic expression of *dprA* prior to competence induction. However, maximal expression of *dprA* results in slightly slower competence shut-off, showing that even ~6,500 DprA molecules per cell were not sufficient for optimal shut-off. Below this point, a sequential loss of competence shut-off was observed, with no shut-off at all observed below ~650 DprA molecules per cell (Figure 6B). As previously reported, the absence of *dprA* delays growth in competent cells 18. Following growth in CEP_*lac*_*-dprA*, *dprA^-^* cells after competence induction revealed an inverse correlation between growth rate and competence shut-off (Figure S2B) with growth more severely affected in conditions where competence was poorly shut-off. This confirmed the importance of competence shut-off for pneumococcal physiology. Taken together, these results show that optimal competence shut-off, but not transformation, requires high cellular levels of DprA.

**Figure 6:**
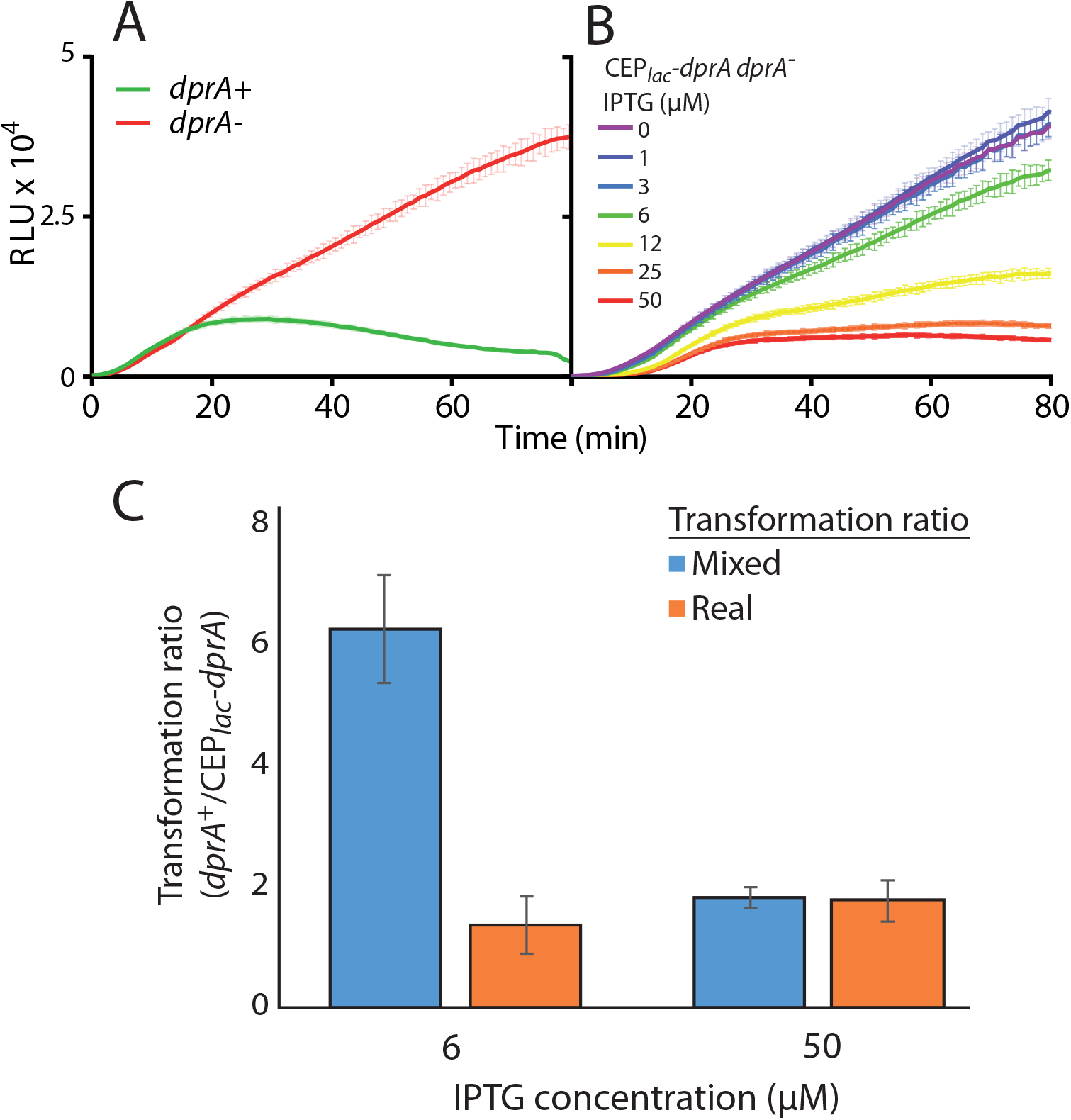
The effect of varying IPTG concentration on competence shut-off in CEP_*lac*_*-dprA*, *dprA^-^*. (A) Competence shut-off tracked using *ssbB-luc* in wildtype (R3584, green) and *dprA^-^* (R3587, red) strains. Results represent averages of triplicate repeats. (B) Competence shut-off tracked using *ssbB-luc* in CEP_*lac*_*-dprA, dprA^-^* (R3834) at varying levels of IPTG. 0 μM, purple; 1 μM, violet; 3 μM, blue; 6 μM, green; 12 μM, yellow; 25 μM, orange; 50 μM, red. Results represent averages of triplicate repeats. (C) Ratios of transformation efficiency between *dprA^+^* (R3584) and CEP_*lac*_*-dprA*, *dprA^-^* (R3833) cells in conditions where CEP_*lac*_*-dprA*, *dprA^-^* transforms optimally but does (50 μM IPTG) or does not (6 μM IPTG) shut-off competence optimally. Comparing real (orange) and mixed (blue) transformation efficiencies allows us to determine the effect of abrogating competence shut-off on transformant fitness (see Figure S5).

### High cellular levels of DprA are important for transformant fitness

To determine if absence of DprA impacts cell viability, colony-forming units (cfu mL^-1^) were compared to OD_492_ readings in competent *dprA^+^* and *dprA^-^* cultures. This revealed a decreased cfu mL^-1^ in *dprA^-^* cultures, strongly suggesting that the lack of competence shut-off affects cell viability as well as negatively impacting growth (Figure S3AB). This was confirmed by tracking competent *dprA^+^* and *dprA^-^* cells by microscopy, where frequent cell lysis was observed in *dprA^-^* cells (Figure S4). The absence of DprA thus impacts cell viability as well as physiology. This suggested that high levels of DprA in competent cells mediate competence shut-off and thus ensure fitness of transformants. To explore this, we took advantage of the fact that a CEP_*lac*_*-dprA*, *dprA^-^* strain grown in 6 μM IPTG transformed at wild-type levels (Figure 3B) while displaying almost no shut-off of competence (Figure 6B). An equal number of competent *dprA^+^* and CEP_*lac*_*-dprA*, *dprA^-^* cells containing different selectable markers (Cm^R^ and Kan^R^ respectively) were mixed and grown in 6 μM IPTG. Cells were then transformed with an *rpsL41* DNA fragment (conferring Sm^R^) and grown together in liquid culture for 3 h 30 min prior to plating and selection. During this time period, *dprA^+^* cells should transform, then rapidly shut-off competence and grow, while CEP_*lac*_*-dprA*, *dprA^-^* cells should also transform but then take longer to shut-off competence, strongly impacting growth (Figure S2). By comparing transformation efficiencies based on the whole mixed population or each individual population, we can directly compare the fitness of transformants (Figure S5). Comparing transformation of individual populations (real transformation ratio) showed that CEP_*lac*_*-dprA*, *dprA^-^* cells transformed at levels comparable to *dprA^+^* cells (Figure 6C). When comparing transformation efficiencies based on the whole mixed population (mixed transformation ratio), *dprA^+^* transformants were recovered over 6-fold more than CEP_*lac*_*-dprA*, *dprA^-^* (Figure 6C), showing that although the individual populations transformed at similar levels, the lack of competence shut-off in CEP_*lac*_*-dprA*, *dprA^-^* resulted in these transformants being out-competed by *dprA^+^* transformants. However, in 50 μM IPTG, where CEP_*lac*_*-dprA*, *dprA^-^* cells transform fully and shut off competence, the *dprA^+^* cells did not out-compete the CEP_*lac*_*-dprA*, *dprA^-^* cells, since both shut-off competence (Figure 6C). This showed that the high levels of DprA produced in competent cells ensure the fitness of transformants by rapidly shutting off competence, promoting the propagation of cells having acquired potentially advantageous genetic alterations.

## Discussion

### Optimal transformation efficiency does not require full *dprA* expression

In this study, we showed that although ~8,300 molecules of DprA are produced per competent cell, only ~650-2,250 are required to ensure optimal transformation efficiency (Figure 3). Below ~650 DprA molecules per cell, a rapid drop in efficiency is observed, although even with only ~40 DprA molecules, transformation still occurs. Despite a ~240-fold decrease in efficiency compared to optimal conditions, this remains high above the ~10,000-fold decrease observed in *dprA^-^* cells 15, 17. Thus, ~40 DprA molecules per cell is enough to carry out transformation, but between ~650 and ~2,250 molecules are required to ensure optimal transformation. This is most likely an over-estimate of the number of DprA molecules required for transformation, as in these conditions, competence shut-off still occurs to some extent (Figure 6B), and thus the majority of available DprA may be mobilized to this end rather than for transformation. Therefore, fewer DprA molecules are probably required for transformation, but this is masked by the secondary role of DprA in competence shut-off.

### Linking the roles of DprA in ssDNA protection and homologous recombination during transformation

By using radiolabeled donor DNA, we were able to demonstrate that the protection of transforming ssDNA paralleled transformation efficiency in competent cells expressing varying levels of DprA. The fact that *dprA^AR^* and *dprA^QNQ^* mutant cells, in which DprA cannot dimerize (and as a result interacts poorly with ssDNA) or interact with RecA, respectively, also do not protect ssDNA, shows that three interactions are necessary for ssDNA protection. Firstly, DprA must be able to form dimers, secondly, DprA dimers must be able to interact with transforming ssDNA and finally, DprA must be able to interact with RecA. Taken together with the fact that both DprA and RecA are crucial for ssDNA protection 17, this result shows that the recombination mediator action of DprA also ensures protection of transforming ssDNA from degradation. These results demonstrate for the first time that the protective role of DprA on ssDNA is mediated directly by the loading of RecA onto ssDNA. This strongly suggests that the polymerization of RecA along transforming ssDNA prevents nuclease access, mediating protection.

### Cellular levels of DprA are high during competence to ensure optimal competence shut-off

The unexpected finding of a key role for DprA in competence shut-off 18 makes this protein unique in playing two central roles in two intrinsically linked processes. These two roles involve starkly different mechanisms and protein domains. DprA forms dimers by self-interaction via the eRF domain (Figure 1B), and during transformation these dimers bind to transforming ssDNA via the DNA-binding domain at the centre of the DprA protein 16. DprA also interacts with RecA via the eRF domain, with a strong overlap between RecA-binding and dimerization interfaces. The role of DprA in competence shut-off involves the N-terminal SAM domain, which binds strongly to ComE~P, the dimeric active form of the ComE response regulator. It was suggested that DprA either interacts with ComE~P to remove it from its target DNA, disrupts the phosphorylated ComE~P dimer or sequesters ComE~P to promote competence shut-off 18. While ~80,000 molecules of ComE are produced during competence 37, 10-fold less DprA molecules are produced 18. However, since DprA primarily interacts with ComE~P, which likely makes up a minority fraction of cellular ComE, DprA may in fact be in excess compared to its ComE~P target. The ratio of ComE/ComE~P was shown to govern competence initiation and shut-off, with ComE~P and ComE competing for access to the same early *com* gene promoters. ComE~P activates these promoters while ComE represses them 37. Interaction of DprA with ComE~P thus alters this ratio in favor of the repressor ComE by removing ComE~P, mediating shut-off.

Here we have shown that optimal competence shut-off requires full expression of *dprA*, with a stark decrease in shut-off observed below 5,000 DprA molecules per competent cell. In light of our findings, antiactivation of ComE~P on its target DNA appears the least likely hypothesis for DprA activity in competence shut-off, since such high levels of DprA are required for optimal shut-off and ComE~P has only nine genetic targets 37. We suggest that DprA more likely either dephosphorylates ComE~P dimers or sequesters ComE~P to prevent access to the early *com* gene promoters. Either of these activities would potentially require high cellular levels of DprA to shift the ComE/ComE~P ratio in favour of ComE and mediate competence shut-off.

Our findings show that high cellular levels of DprA are key for the shut-off of pneumococcal competence but not necessary for the primary role of DprA as a RMP in transformation. The secondary role of DprA in competence shut-off is an acquired role 18. Competence is controlled by ComDE in *mitis* and *anginosus* groups of streptococci but by ComRS in the remaining streptococci 18. Comparing the evolution rates of DprA among streptococci showed that the SAM domain of DprA, which interacts with ComE~P to shut-off competence, shows a higher rate of evolution than the eRF domain in streptococci regulating competence via ComDE but not those using ComRS. This strongly suggested that the control of competence shut-off by DprA is conserved among the *anginosus* and *mitis* groups of streptococci 18. In light of our findings, we suggest that the role of DprA in competence shut-off via interaction with ComE~P may be coupled to the high level of cellular DprA necessary for the role of this protein in competence shut-off. This coupling may be conserved among these two groups of streptococci to facilitate the shut-off of competence in species which depend on ComDE for competence regulation.

### High cellular levels of DprA ensure transformant fitness to maximise adaptive potential

The lack of DprA in competent cells was previously shown to be detrimental for pneumococcal physiology *in vitro* 18 as well as bacteraemia and pneumonia *in vivo* 24. Here, we have shown that a competence induces cell lysis in a *dprA^-^* population (Figure S3). Furthermore, high cellular levels of DprA during competence are crucial for the fitness of transformants (Figure 6C). These findings demonstrate that although low levels of DprA in competent cells can ensure optimal transformation (Figure 3), high cellular levels of DprA are required to ensure the propagation and survival of these transformants. The pneumococcus colonizes the human nasopharynx, an environment it shares with many other bacteria 39. In a situation where competence is induced in response to a stress such as expose to antibiotics 40, the whole pneumococcal population becomes competent, but only a minority may gain a selective advantage allowing survival in response to the initial stress. This minority, despite having a selective advantage over the rest of the pneumococcal population, would be out-competed by neighbouring species which may already be resistant to the stress, and the pneumococcal population would not survive. However, competence is not always a reactive mechanism. For example, capsule switching by transformation, which allows vaccine escape 41, occurs spontaneously within mixed populations and not as a response to vaccine administration 42. In this situation, high cellular levels of DprA allow capsule switch transformants to compete with the rest of the niche population in the absence of selective pressure. Thus, by ensuring rapid and efficient competence shut-off, high cellular levels of DprA ensure the fitness of pneumococcal transformants, allowing them to propagate and survive. This in turn maximises the adaptive potential of the pneumococcus (Figure 7).

**Figure 7:**
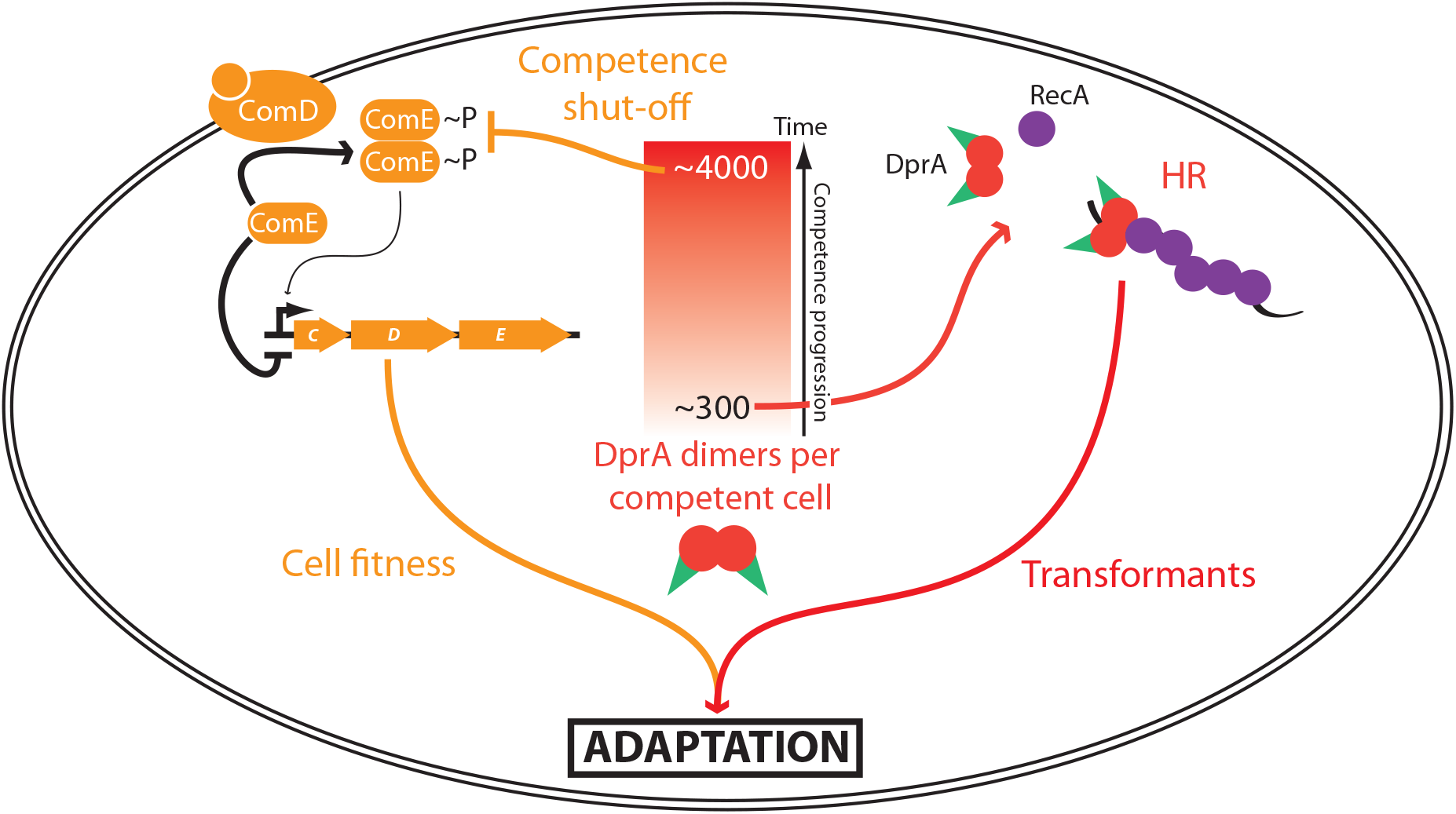
Cellular levels of DprA during competence dictate transformant fitness and adaptive potential. The number of DprA dimers present in competent cells evolves over time during the induction of competence (Red gradient box). Full expression of DprA (~4,000 dimers) is required for competence shut-off (orange). DprA dimers either dephosphorylate or sequester ComE~P dimers, which shifts the ratio of ComE/ComE~P in favour of the repressive ComE, promoting competence shut-off. However, only ~300 DprA dimers are required to ensure optimal transformation efficiency (red). DprA is necessary to generate transformants, and to ensure cell fitness by shutting off competence, ensuring the adaptive potential of transformation can be realized.

### Concluding Remarks

The induction of pneumococcal competence is crucial to the lifestyle of the pneumococcus, since it mediates transformation and thus allows adaptation. However, uncontrolled competence induction is detrimental to pneumococcal physiology, affecting both pneumococcal physiology *in vitro* 18 and survival *in vitro* (Figure S3) and *in vivo* 24. Since DprA interacts directly with ComE~P to mediate the shut-off of pneumococcal competence, there is a direct link between DprA and pneumococcal fitness. It has been suggested that the link between competence and fitness provides a selective pressure for maintenance of an intact *com* regulon, which in turn provides the adaptive benefit of transformation 43. Here, we have shown that the high cellular levels of DprA are crucial for optimal competence shut-off and for the fitness of transformants. We suggest that these high levels have evolved to ensure tight control of competence shut-off, providing a fitness advantage to pneumococcal transformants and maximizing the adaptive potential that transformation provides. Since DprA is required both to generate pneumococcal transformants and ensure their fitness, it emerges as an interesting therapeutic target to drastically reduce the adaptive potential of this human pathogen in response to treatment.

## Materials and Methods

### Bacterial strains, primers and transformation

The pneumococcal strains and primers used in this study can be found in Table S1. Standard procedures for transformation and growth media were used 44. In this study, cells were rendered unable to spontaneously develop competence by replacing the *comC* gene which encodes CSP1 with an allelic variant encoding CSP2 45, since ComD1 is unable to respond to CSP2 46. Unless described, pre-competent cultures were prepared by growing cells to a OD_550_ of 0.1 in C+Y medium before 10-fold concentration and storage at −80°C as 100 μL aliquots. Antibiotic concentrations (μg mL^-1^) used for the selection of *S. pneumoniae* transformants were: chloramphenicol (Cm), 4.5; kanamycin (Kan), 250; spectinomycin (Spc), 100; streptomycin (Sm), 200. For the monitoring of growth and *luc* expression, precultures were gently thawed and aliquots were inoculated (1 in 100) in luciferin-containing 34 C+Y medium and distributed (300 ml per well) into a 96-well white microplate with clear bottom. Relative luminescence unit (RLU) and OD values were recorded throughout incubation at 37°C in a Varioskan luminometer (ThermoFisher). The *ssbB-luc* reporter gene was transferred from R895 as previously described 38. The *dprA*::*spc^21^*^C^ cassette was transferred from R1800 as previously described 18.

### Assessing transformation efficiency of CEP_*lac*_*-dprA, dprA^-^*

To determine the transformation efficiency of R3834 (CEP_*lac*_*-dprA, dprA^-^*), we used a previously-described transformation protocol 44. Briefly, pre-competent cells were prepared by growth to OD550 0.1 in 3 mL C+Y medium containing varying concentrations of IPTG (1, 3, 6, 12, 25, 50 μM), centrifuged to harvest cells and resuspended in 300 μL C+Y with 15% glycerol before being split into 100 μL aliquots containing ~10^7^ cells. For transformation, 100 μL aliquots of pre-competent cells were resuspended in 900 μL fresh C+Y medium with 100 ng mL^-1^ CSP and appropriate IPTG concentrations and incubated at 37°C for 10 minutes. Transforming DNA was then added to a 100 μL aliquot of this culture, followed by incubation at 30°C for 20 minutes. Cells were then diluted and plated on 10 mL CAT agar with 5% horse blood and appropriate concentrations of IPTG before incubation at 37°C for 2 h. A second 10 mL layer of CAT agar with streptomycin (200 μg mL^-1^) was added to plates to select transformants, and plates without antibiotic were used as comparison to calculate transformation efficiency. Plates were incubated overnight at 37°C. Transforming DNA was either 25 ng of R304 47 genomic DNA or 6 × 10^−3^ ng (equivalent of ~3 × 10^6^ DNA molecules) of a 1,982 bp PCR fragment amplified with primer pair rpsL3-rpsL4. Since this was transformed into ~10^7^ competent cells, this represented roughly 1 DNA molecule per 3 competent cells. Both R304 genomic DNA and the PCR fragment transferred the *rpsL41* point mutation, conferring streptomycin resistance. Results presented are averages of triplicate repeats. Statistical analyses used were unpaired t-tests done using GraphPad Prism 7.04.

### Assessing competence shut-off and growth of CEP_*lac*_*-dprA, dprA^-^, ssbB-luc*

To determine the effect of varying cellular DprA levels on competence shut-off, we used a previously-described protocol 34, with modifications. Briefly, cells were grown to OD_550_ 0.2 in 2 mL C+Y medium containing varying concentrations of IPTG (1, 3, 6, 12, 25, 50 μM), harvested and resuspended in 1 mL C+Y with 15% glycerol before being split into 100 μL aliquots. Cells were then diluted 100-fold in 270 μL C+Y with 0.5 mM luciferin and appropriate IPTG concentration in a 96-well plate before incubation at 37°C in a Varioskan luminometer (ThermoFisher) with luminometry and OD readings every 10 minutes. When an OD_492_ of ~0.05 was reached, 200 ng mL^-1^ CSP was added to each well, and the plate was incubated for 37°C with luminometry readings every 2 minutes for 80 minutes to track the shut-off of competence. Results presented are averages of triplicate repeats. To assess the effect of varying cellular DprA levels on growth, a similar experiment was carried out, although no luminometry readings were taken, and optical density was measured every 10 minutes for 3 hours.

### Co-transformation analysis of transformant fitness

Pre-competent *dprA^+^* (R3584, Cm^R^) and CEP_*lac*_*-dprA*, *dprA^-^* (R3833, Kan^R^, Spc^R^) cells were grown in varying concentrations of IPTG (50, 6 μM) and stored at - 80°C as 50 μL aliquots. *dprA^+^* cells were then mixed 1:1 with CEP_*lac*_*-dprA*, *dprA^-^* cells and diluted 10-fold in C+Y containing either 50 or 6 μM IPTG and competence was induced by addition of 200 ng mL^-1^ CSP. Cells were then incubated for 10 minutes at 37°C before addition of 100 ng mL^-1^ transforming DNA (PCR fragment of 1.4 kb possessing *rpsL41* point mutation conferring streptomycin resistance, Sm^R^) before further incubation at 30°C for 20 minutes. 1.4 mL fresh C+Y was added to cultures, followed by incubation at 37°C for a further 3 h 30 min. Cultures were then diluted and cells were plated on CAT agar supplemented with 5% horse blood and appropriate antibiotics. Cells were plated to determine total cells present (no selection), *dprA^+^* cells present (Cm^R^), CEP_*lac*_*-dprA*, *dprA^-^* cells present (Kan^R^), *dprA^+^* transformants (Cm^R^, Sm^R^) and CEP_*lac*_*-dprA*, *dprA^-^* transformants (Kan^R^, Sm^R^).

## Acknowledgements

We thank Nathalie Campo and Mathieu Bergé for critical reading of the manuscript. We also thank Mathieu Bergé for the *comC2D1* construct. We thank Bernard Martin for help initiating the project and for useful discussions. We thank Anne-Lise Soulet for experimental assistance. This work was funded by the Centre National de la Recherche Scientifique, University Paul Sabatier, Agence Nationale de la Recherche (grants ANR-10BLAN-1331 and EXStasis-17-CE13-0031-01).

## Supplementary Figure Legends

**Figure S1: Validating the CEP_*lac*_ expression system for use in *S. pneumoniae*.** (A) Genetic organization of CEP_*lac*_-luc expression platform. Organization as in Figure 2A but with *dprA* replaced by *luc.* (B) Expression profile of CEP_*lac*_-luc in different concentrations of IPTG. Results represent averages of triplicate repeats. To simplify the figure, a single growth curve is presented to represent all growth curves, which were identical. (C) Maximum expression levels of CEP_*lac*_-luc expressed with different concentrations of IPTG. Results represent averages of triplicate repeats. Strain used: CEP_*lac*_-luc, R3310.

**Figure S2: The impact of competence on growth when varying cellular levels of DprA are present.** (A) Growth of *dprA^+^* (R3584) and *dprA^-^* (R3587) cells after CSP addition at 0 min. (B) Growth of CEP_*lac*_*-dprA*, dprA^-^ (R3833) cells in varying concentrations of IPTG after CSP addition at 0 min. (C) Growth of *dprA^+^* (R3584) and *dprA^-^* (R3587) cells in absence of CSP. (D) Growth of CEP_*lac*_*-dprA*, dprA^-^ (R3833) cells in varying concentrations of IPTG in absence of CSP.

**Figure S3: The effect of the absence of *dprA* on survival of competent cells.** (A) Viable cells in *dprA^+^* (R3584, blue) and *dprA^-^* (R3587, orange) cultures at 30 min timepoints after induction of competence by CSP addition. (B) Competence induction (filled lines) and growth (dotted lines) of *dprA^+^* (R3584, blue) and *dprA^-^* (R3587, orange) cells at 15 min time points after induction of competence by CSP addition.

**Figure S4: Observing viability of competent cells lacking DprA.** (A) Images of competent *dprA^+^* (R3584) cells taken every 20 minutes from 30 minutes after competence induction. Red arrow indicates estimated duration of competence window in the population, based on Figure S3B. Green arrow indicates estimated time when exponential growth resumes post-competence. (B) Images of competent *dprA^-^* (R3587) cells taken every 20 minutes from 30 minutes after competence induction. Red arrow indicates estimated duration of competence window in the population, since excess of CSP is present in C+Y agar during observation, resulting in permanent cycling of competence. Three images taken to represent *dprA^-^* cultures, where some cells appear go grow while others lyse. Yellow arrows, pre-lysis cells; blue arrows, post-lysis cell remnants.

**Figure S5: Explanatory diagram of the experiment carried out in Figure 7.** An equal cellular density of pre-competent *dprA^+^* (R3584, Cm^R^) and CEP_*lac*_*-dprA*, *dprA^-^* (R3833, Kan^R^) cells was mixed before being induced to competence by CSP, exposed to transforming DNA (*rpsL41* PCR, conferring Sm^R^) and diluted to allow phenotypic expression. After phenotypic expression, cells were diluted and plated to select for either all cells (no selection), *dprA^+^* cells (Cm^R^), CEP_*lac*_*-dprA*, *dprA^-^* cells (Kan^R^), *dprA^+^* transformants (Cm^R^, Sm^R^) and CEP_*lac*_*-dprA*, *dprA^-^* transformants (Kan^R^, Sm^R^). From these values, the real and mixed transformation efficiencies were calculated for each population, and real and mixed transformation ratios were determined. Comparing the ratios in conditions where competence shut-off is present (50 μM IPTG) or absent (6μM IPTG) allows us to define the effect of competence shut-off on transformant fitness.

